# Maximizing Quantitative Phosphoproteomics of Kinase Signaling Expands the Mec1 and Tel1 Networks

**DOI:** 10.1101/2020.03.25.008367

**Authors:** Vitor Marcel Faca, Ethan Sanford, Jennifer Tieu, Shannon Marshall, William Comstock, Marcus Smolka

**Affiliations:** Department of Molecular Biology and Genetics, Weill Institute for Cell and Molecular Biology, Cornell University, Ithaca, NY 14853, USA; Department of Biochemistry and Immunology and Cell-Based Therapy Center, Ribeirao Preto Medical School, University of Sao Paulo, Ribeirao Preto, SP, 14049-900, Brazil

**Author notes:** Correspondence and requests for materials should be addressed to M.B.S. These authors contributed equally to this work (Vitor Marcel Faca, Ethan Sanford).

**Keywords:** Phosphoproteome, SILAC, Kinase, HILIC, Mec1, Tel1, ATR

## Abstract

Global phosphoproteome analysis is crucial for comprehensive and unbiased investigation of kinase-mediated signaling. However, since each phosphopeptide represents a unique entity for defining identity, site-localization, and quantitative changes, phosphoproteomics often suffers from lack of redundancy and statistical power for generating high confidence datasets. Here we developed a phosphoproteomic approach in which data consistency among experiments using reciprocal stable isotope labeling defines a central filtering rule for achieving reliability in phosphopeptide identification and quantitation. We find that most experimental error or biological variation in phosphopeptide quantitation does not revert in quantitation once light and heavy media are swapped between two experimental conditions. Exclusion of non-reverting data-points from the dataset not only reduces quantitation error and variation, but also drastically reduces false positive identifications. Application of our approach in combination with extensive fractionation of phosphopeptides by HILIC identifies new substrates of the Mec1 and Tel1 kinases, expanding our understanding of the DNA damage signaling network regulated by these kinases. Overall, the proposed quantitative phosphoproteomic approach should be generally applicable for investigating kinase signaling networks with high confidence and depth.

## INTRODUCTION

Protein phosphorylation is of central importance in both normal physiology and pathological conditions. Phosphorylation-mediated switches regulated by protein kinases and protein phosphatases can affect protein structure and function, with consequences in enzymatic activity, protein localization, protein interactions and turnover [1]–[3]. Understanding the in vivo action of kinases requires unbiased mapping of the specific phospho-signaling events they mediate, which is achieved mainly using mass-spectrometry based approaches. Both instrumentation and bioinformatic tools applied for phosphopeptide identification have been continuously evolving [4]–[6], culminating in larger phosphoproteomic datasets in recent years [7]–[12]. In addition to in depth coverage of the phosphoproteome, the gold standard for comprehensive mapping of kinase-mediated signaling also requires quantitative analysis of each phosphopeptide or phosphorylation site to monitor its abundance in conditions of active kinase compared to conditions in which the kinase of interest is chemically and/or genetically ablated [13]–[15]. Various quantitative mass spectrometric approaches have been applied for the identification of kinase substrates, including stable isotope labeling in cell culture (SILAC) [15]–[17] and various isobaric labelling strategies such as tandem mass tag (TMT) [18], [19]. In a recent systematic comparison of quantitative phosphoproteomic strategies, SILAC was considered the most accurate, although TMT-based analyses yielded better coverage of the phosphoproteome [6]. SILAC is based on peptide precursor ion quantification to detect and quantify, in relative terms, the ratio between heavy isotope-labeled or unlabeled (light) amino acids incorporated metabolically into cells [20], [21]. Such an approach allows early mixing of labeled protein extracts in phosphoproteomic workflows to minimize technical variation.

Despite the broad use of mass spectrometry for investigating kinase-mediated phosphorylation signaling responses, major challenges remain for systematically achieving high confidence identification and quantitation analysis of phosphopeptides. The key difference between proteomics and phosphoproteomics is that when analyzing protein abundance and/or interactions (as in affinity-purification mass spectrometry), the analysis is based on identification and quantification of multiple redundant representative peptides for a given protein, whereas in phosphoproteomics, phosphopeptides are often unique (non-redundant) species represented by one or a few peptide spectral matches (PSMs) in the dataset,. The lack of multiple redundant events for informing identification, quantification and phospho-site localization hobbles the acquisition of high-quality data due to the low numbers of PSMs per phosphopeptide. The ability of acquiring high quality identification and quantification data is further complicated by the fact that many key phosphopeptides of biological interest are present at very low levels in the pool of phosphopeptides enriched from whole cell lysates. Even in cases when identification of a phosphopeptide based on one or two PSMs is successful, the associated quantitative information can suffer from signal interference derived from sample complexity and other intrinsic technical noise [22]–[25]. As a result, a significant part of the generated phosphoproteomic data is not suited for reliable quantitative analysis and biological inference, representing one of the major bottlenecks in large-scale quantitative phosphoproteomic analysis of kinase-mediated signaling.

Here we report a phosphoproteomic approach for increasing reliability in phosphopeptide identification and quantification, while minimizing loss of data from phosphopeptides with low PSM counts. The approach relies on quantitation consistency among reversed isotopically labeled samples as a filtering step for removing false positive identifications and erroneous quantifications. We find that most experimental error or biological variation in phosphopeptide quantitation does not exhibit reverted quantitation values once light and heavy media are swapped in control experiments. This allows the systematic exclusion of non-reverting data-points from the dataset to reduce not only quantitation error and variation, but also to reduce false positive identifications. The reported approach balances both sensitivity and specificity to detect phosphorylation changes with high confidence, even in the case of phosphopeptides with low PSM counts. Application of our approach identifies new substrates of the yeast Mec1 and Tel1 DNA Damage Response (DDR) kinases, expanding our understanding of the DNA damage signaling network regulated by these kinases. Overall, this simple approach should help enhance the reliability of quantitative phosphoproteomics in biological interrogations of kinase-mediated signaling networks.

## EXPERIMENTAL PROCEDURES

### Yeast cell culture and manipulation

A list of yeast strains used in this study is found in supplemental Table S4. The strain background for all yeast used was S288C. We performed whole ORF deletions of Mec1 and Tel1 kinases using established PCR-based methods for amplifying resistance cassettes containing homology to the target gene. Gene manipulations were verified by PCR. Primera used for gene deletions are available upon request. Yeast were grown at 30°C in synthetic SILAC media lacking arginine and lysine and supplemented with “light” lysine and arginine (^12^C and ^14^N) or supplemented with “heavy” lysine and arginine (L-Lysine ^13^C_6_,^15^N_2_.HCl and L-Arginine ^13^C_6_,^15^N_4_.HCl). Media was also supplemented with excess L-proline to prevent conversion of arginine to proline.

### Sample preparation for Phosphoproteomic Analysis

200-300mL of yeast was grown in either “heavy” or “light” SILAC media to mid-log phase and treated as described in the figure legend and the text, depending on the experiment. Cells were pelleted at 1000xg and washed once with TE (10mM Tris pH 8.0, 5mM EDTA) buffer containing 1mM PMSF. Cells were lysed by bead beating with 0.5mm glass beads for 3 cycles of 10 minutes with 1-minute rest time between cycles at 4°C in lysis buffer (150mM NaCl, 50mM Tris pH 8.0, 5mM EDTA, 0.2% Tergitol type NP40) supplemented with protease inhibitor cocktail (Pierce), 5 mM sodium fluoride and 10 mM β-glycerophosphate. 5-7mg of each light and heavy labeled protein lysate was denatured and reduced with 1% SDS and 5mM DTT at 42°C, then alkylated with 25mM iodoacetamide. Lysates (light and heavy) were mixed and precipitated with a cold solution of 50% acetone, 49.9% ethanol, 0.1% acetic acid. Post-precipitation protein pellet was then resuspended in 2M urea and subsequently digested with TPCK-treated trypsin overnight at 37°C. Phosphoenrichment was performed using a High-Select™ Fe-NTA phosphopeptide enrichment kit (ThermoFisher Scientific, cat# A32992) as described in the manufacturer’s instructions. Purified phosphopeptides were then dried in a SpeedVac and fractionated via HILIC chromatography as described below.

### HILIC fractionation

Dried phosphopeptide samples were reconstituted in 15 μL H_2_O, 10 μL 10% formic acid (v/v), and 60 μL HPLC-grade acetonitrile. 80 μL of the reconstituted sample was injected and fractionated by hydrophilic interaction liquid chromatography (HILIC) using a TSK gel Amide-80 column (2 mm x 150 mm, 5 μm; Tosoh Bioscience). Three solvents were used for the gradient: buffer A (90% acetonitrile), buffer B (75% acetonitrile and 0.005% trifluoroacetic acid), and buffer C (0.025% trifluoroacetic acid). A short gradient was used for the mock control and Mec1 experiments and consisted of 100% buffer A at time = 0 min; 88% of buffer B and 12% of buffer C at time = 5 min; 60% of buffer B and 40% of buffer C at time = 30 min; and 5% of buffer B and 95 % of buffer C from time = 35 to 45 min in a flow of 150 µl/min. 30-second fractions were collected between 9 and 18 minutes. An extended gradient was used for the Tel1 experiment and consisted of 100% buffer A at time = 0 min; 94% buffer B and 6% buffer C at t = 3 min; 65.6% buffer B and 34.4% buffer C at t = 30 min with a curve factor of 7; 5% buffer B and 95% buffer C at t = 32 min; isocratic hold until t = 37 min; 100% buffer A at t = 39 to 51 min at a flow rate of 150 μL/min. One-minute fractions were collected between minutes 8 and 10 of the gradient; 30-second fractions between minutes 10 and 26; and two-minute fractions between minutes 26 and 38 for a total of 40 fractions. Individual fractions were dried in speedvac and submitted to LC-MS/MS analysis.

### Phosphoproteomics data acquisition

Individual phosphopeptide fractions were resuspended in 0.1% trifluoroacetic acid and subjected to LC-MS/MS analysis in an UltiMate™ 3000 RSLC nano chromatographic system coupled to a Q-Exactive HF mass spectrometer (Thermo Fisher Scientific). The chromatographic separation was carried out in 35-cm-long 100-µm inner diameter column packed in-house with 3 µm C_18_ reversed-phase resin (Reprosil Pur C18AQ 3μm). Q-Exactive HF was operated in data-dependent mode with survey scans acquired in the Orbitrap mass analyzer over the range of 380 to 1800 m/z with a mass resolution of 60,000 (at m/z 200). MS/MS spectra was performed selecting the top 15 most abundant +2, +3 or +4 ions and a with an precursor isolation window of 2.0 m/z. Selected ions were fragmented by Higher-energy Collisional Dissociation (HCD) with normalized collision energies of 28 and the mass spectra acquired in the Orbitrap mass analyzer with a mass resolution of 15,000 (at m/z 200), AGC target set to 1e^5^ and max injection time set to 120ms. A dynamic exclusion window was set for 30 seconds.

### Phosphopeptide and phosphosite identification

The peptide identification and quantification pipeline relied on TPP tools [26]. The search engine used was Comet (v. 2019.01.1) [27]. Search parameters included semi-tryptic requirement, 20ppm for the precursor match tolerance, differential mass modification of 8.0142 for lysine, 10.00827 for arginine, 79.966331 for phosphorylation of serine, threonine and tyrosine, 987.976898 for phosphorylation dehydration, and static mass modification of 57.021465 for alkylated cysteine residues. The protein sequence database was the SGD yeast supplemented with the decoy reversed sequences and common contaminants (downloaded in Aug 2019, 11968 entries). Original ThermoScientific .raw files were converted to mzXML before the search with Comet. After searches, peptides were filtered and scored using PeptideProphet algorithm [28]. Additional options for scoring were: minimum probability of 0.5, minimum peptide length of 7 amino acid residues, accurate mass binning, use Phospho information and restricted to 2, +3 and +5 ion charge states. After scoring and filtering, relative quantitation based on SILAC were obtained using Xpress and specific parameters were: mass tolerance of 0.005 daltons; minimum number of chromatogram points needed for quantitation = 1; number of isotopic peaks = 0. Phosphopeptides were then evaluated by PTMProphet [29] in order to obtain accurate phosphosite localization score. The complete lists of identified, quantified, scored and filtered phosphopeptides were further processed using a R-script developed in-house. The script separates phosphosites with high PTMProphet probability (>0.9) from those with ambiguous localization containing 2 or more adjacent potentially phosphorylated residues, here denominated “clusters”. Separately, high confidence phosphosites and clustered phosphosites had their SILAC quantitation median calculated and additional R-scripts were used for combining, correlating and plotting the data.

## RESULTS

### Error and Variation in SILAC-based Phosphopeptide Quantitation is Unidirectionally Biased

We set out to develop an approach to maximize confidence in quantitative data from phosphoproteomic experiments. We postulated that SILAC-based quantitation might be particularly well suited for separating meaningful biological changes from: (1) aberrant quantitation during data processing (herein referred as “Error”), and/or (2) changes in phosphopeptide abundance unintentionally introduced during sample handling (herein referred as “Variation”). If both <u>E</u>rror and/or <u>V</u>ariation (EV) are mostly associated with artefacts that are independent of true biological differences in the cell lines or drug treatment conditions being compared, phosphoproteomic analysis should reveal a unidirectional bias in the generated ratios of data points reflecting EVs (Fig. 1A-C). We further reasoned that a strong bias in EVs would enable their systematic exclusion from large-scale phosphoproteomic datasets and, in principle, enable the generation of high confidence quantitative data even from phosphopeptides with only one PSM detected in each reciprocal, labeling swapping SILAC experiment.

**Figure 1.**
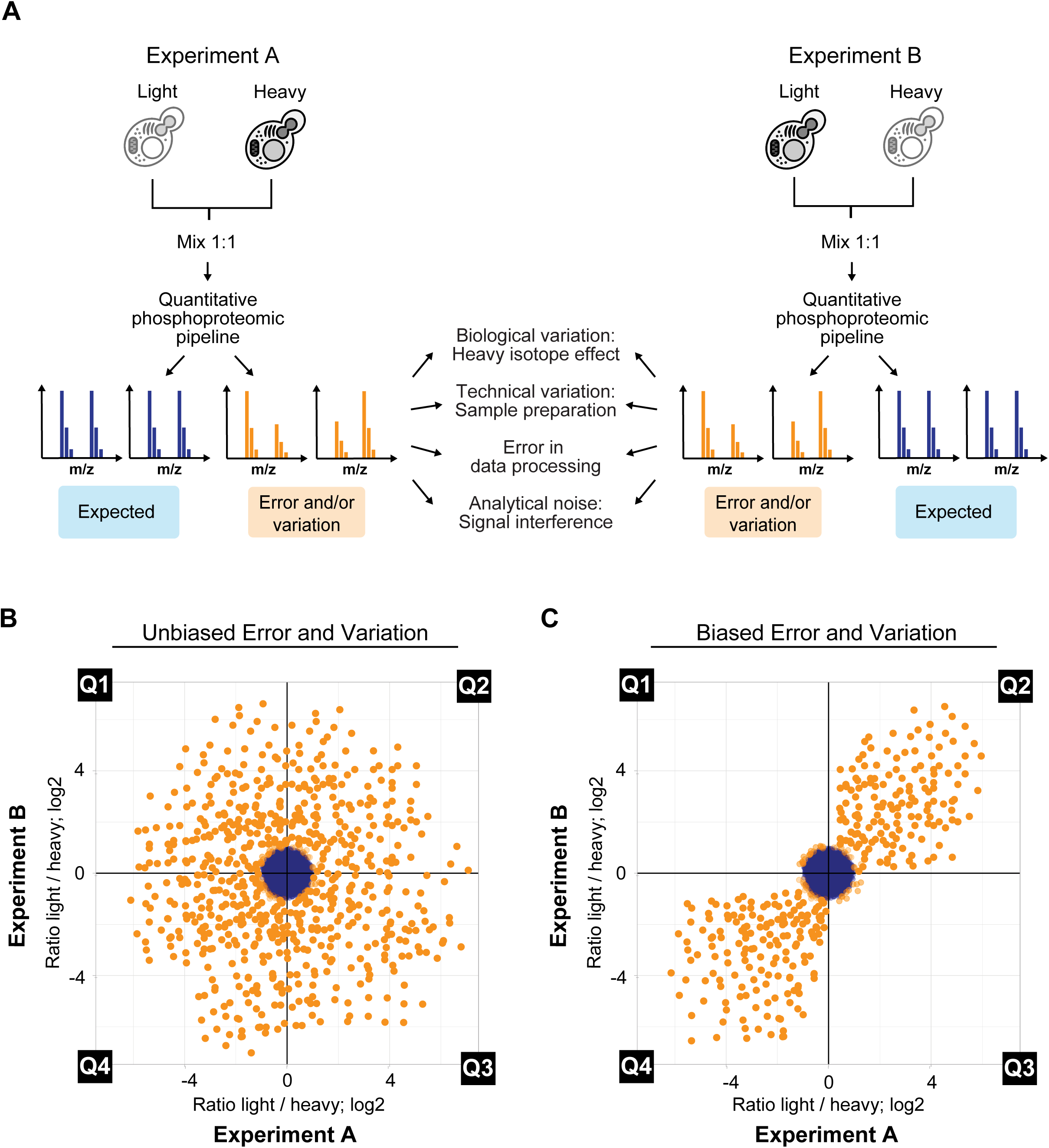
Modeling Outcomes of SILAC Reciprocal Labeling as a Means to Reduce Technical Error and/or Variation (EV). A. Workflow showing reciprocal labeling scheme with a forward experiment (Experiment A, left) and a reverse experiment (Experiment B, right) with anticipated outcomes and proposed causes of EVs (middle). B. Anticipated distribution of false positives in a comparison of two identical samples if error and variation occurred randomly and independent of isotopic labeling. C. Anticipated distribution of false positives in a comparison of two identical samples if error and variation were unidirectionally biased (i.e. similar ratio in both a forward and reverse experiment).

To test this idea, we mixed equal amounts of protein extracts from budding yeast grown in light (^12^C^14^N arginine and lysine) or heavy (^13^C^15^N arginine and lysine) SILAC media (see Materials and Methods for experimental details) and subjected lysates to a quantitative phosphoproteomic pipeline and data analysis described in Figure S1. An independent biological replicate was performed to mimic a reciprocal, labeling swapping experiment. As shown in Figure 1A, data points with a SILAC ratio not reflecting the expected 1:1 ratio (simulated within a 33% coefficient of variation, or approximately a 2-fold change), were considered to reflect methodological error and/or variation. Comparison of experiments A and B (control and reciprocal label swap) should reveal if the error and/or variation exhibit any biased distribution in a quantitative plot (Figs. 1B-C). As shown in Fig. 2A (see Table S1 for detailed dataset), separate experiments revealed thousands of data points outside a simulated range of 33% coefficient of variation (indicated in yellow). We reasoned that these points reflect EVs in the experiment. Notably, comparison of the ratio of each phosphopeptide in experiments A and B revealed a clear bias in EV distribution toward quadrants Q2 and Q4 (Figs. 2B-C) such that 92% of all EVs fell within these quadrants. Notably, EVs accounted for 17% of all phosphopeptides present in our dataset when considering phosphopeptides with 1 PSM in each experiment, underscoring the importance of their exclusion. Data points in Q2 and Q4 represent phosphopeptides whose SILAC ratios did not revert in the reciprocal experiment. Overall, these results reveal that the use of a SILAC labeling swap in phosphoproteomic experiments allows efficient detection of intrinsic EV in the dataset, which may be used for achieving high confidence quantitative analysis, even for phosphopeptides represented by a low number of PSMs. This ability to filter signal from noise, even when PSM numbers are low, is crucial for phosphoproteomic experiments which often rely on difficult-to-detect phosphopeptides. In fact, approximately a third of the data points in the correlation plot shown in Fig. 2B reflect phosphopeptides with only one PSM in one of the experiments (see Table S1).

**Figure 2.**
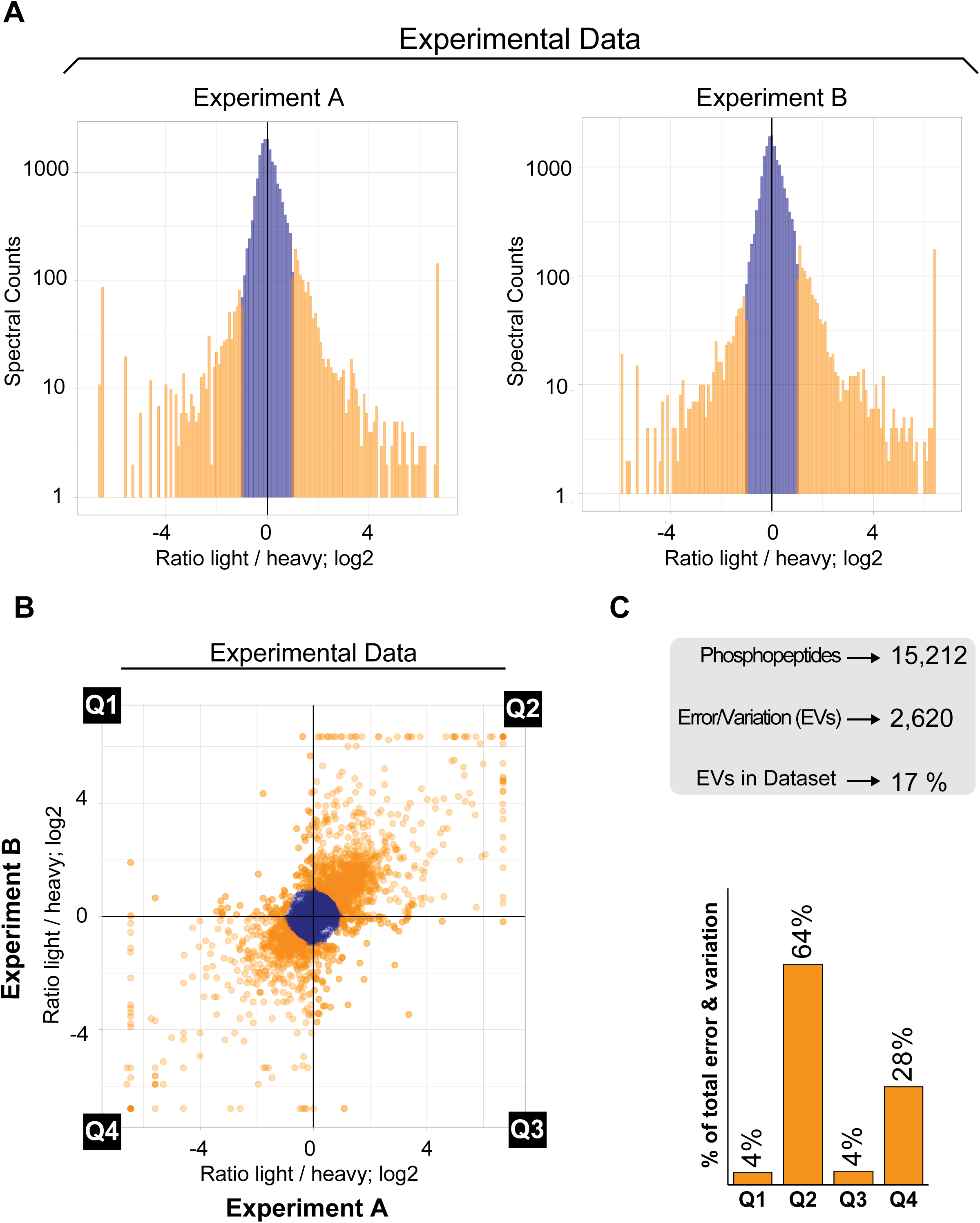
Reciprocal Labeling in an Isogenic Yeast Cell Line Reveals Extensive Error and Variation that is Unidirectionally Biased. A. Histograms for SILAC ratios of two independent phosphoproteome experiments comparing isogenic wild-type *S. cerevisiae* as depicted in Figure 1A. EVs are colored in orange. B. Scatterplot comparing experimental data from the two SILAC experiments shown in A. EVs are colored in orange. C. Histogram showing unidirectional bias of error and variation toward quadrants 2 and 4 in plot from B.

### Data Filtering Approaches for Reducing Error and Variation

To apply data filtering approaches for efficiently eliminating EVs while minimizing loss of data, we evaluated the effects of imposing thresholds on the minimal number of observations (PSMs) required for each phosphorylation site identified. While each phosphosite requires at least 2 observations (1 in each of the reciprocal experiments) to be shown in the correlation plot, increasing the requirement for 2 or more observations in each experiment decreased the proportion of EVs in relation to the entire dataset (Fig. 3A). Considering specifically data points present in Q1 and Q3, where inverse correlation is expected between phosphopeptide ratios in reciprocal experiments, we find that the proportion of EVs is about 1.5% when considering 1 or more PSM in each experiment. This EV proportion is reduced by approximately half, to 0.8% of the data points, when a minimum of 2 observations is required in each experiment (Fig. 3B). However, this additional requirement also decreased sensitivity, reducing the total number of data points from 15,212 to 10,498 (Table S1).

**Figure 3.**
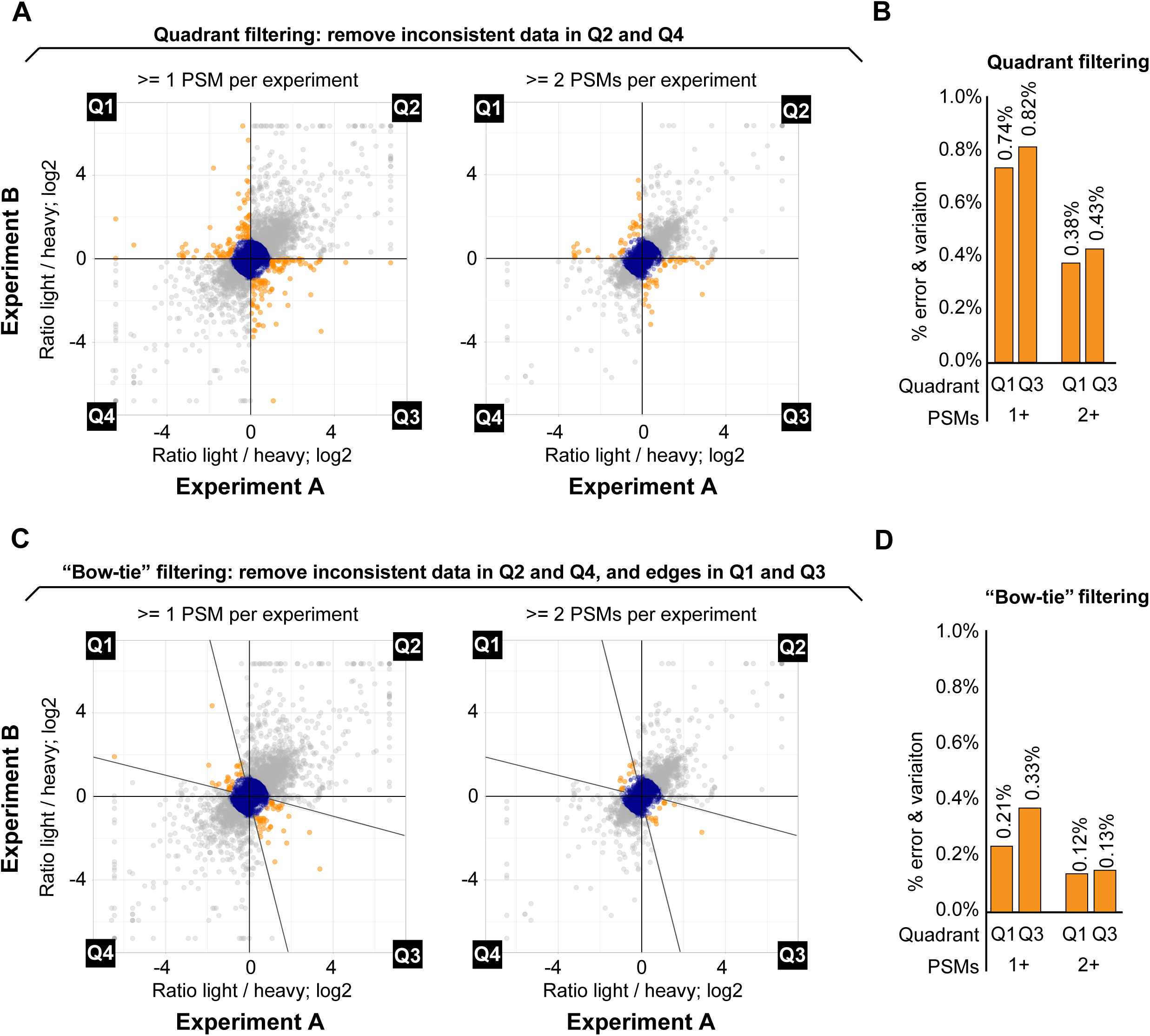
Data Filtering Based on Quantitation Consistency Drastically Reduces Error and Variation. A. Scatterplot from Figure 2B. indicating the “Quadrant” filtering scheme (gray data points removed in Q2 and Q4) and additional filtering based on the requirement for at least 2 PSMs per experiment for each data point (plot on the right). B. Histogram showing EVs in Q1 and Q3 (orange data points) from A as a percentage of total dataset using either 1 PSM or 2 PSM filtering. C. Scatterplot from Figure 2B. indicating the “Bow-tie” filtering scheme (gray data points removed) and additional filtering based on the requirement for at least 2 PSMs per experiment for each data point (plot on the right). For Bow-tie filtering, in addition to removing EVs in Q2 and Q4, data points in Q1 and Q3 were required to be within an interval of correlation correspondent to 4-fold of the log2 scale. D. Histogram showing highlighted points in quadrants 1 and 3 from C as a percentage of total dataset using either 1 PSM or 2 PSM filtering.

During our EV analyses, we noticed a clear prevalence of data points close to the X-axis and Y-axis in Q1 and Q3 (Figs. 3A and 3C), revealing data points with a deviated ratio in only one of the experiments. By employing a simple “quadrant filtering” approach, whereby points in Q2 and Q4 are excluded, and points in Q1 and Q3 are kept, we cannot exclude highly variable phosphopeptide measurements that are also likely the result of error and/or variation (Fig. 3C). To circumvent this issue and more efficiently remove EVs for improved data quality, we designed an alternative filtering approach where data points in Q1 and Q3 were required to be within an interval of correlation correspondent to 4-fold of the log2 scale (herein referred to “Bow-tie filtering”) (Figs. 3C). As shown in Figures 3C-D, the use of Bow-tie filtering, even where peptides with 1 PSM in each experiment were included, reduced the proportion of EVs to 0.54% of the dataset. When Bow-tie filtering was combined with the threshold of at least 2 PSMs per experiment, the proportion of EVs again dropped by approximately half to 0.25% of the dataset. These results reveal that the ability to identify EVs in SILAC-based phosphoproteomic experiments allows the utilization of filtering strategies that drastically reduce error and variation in the dataset, therefore increasing the confidence in the data even when considering phosphopeptides represented by a single PSM per experiment.

### Eliminating Error and Variation in Quantitation Reduces Decoy Identifications

SILAC labeling with stable isotopes shifts the mass of parent ions and their fragments in both MS1 and MS2, respectively. We reasoned that this mass shift should enable more efficient exclusion of false positive identifications in the dataset, since a misidentification would need to occur in both reciprocal experiments and be consistent between two parental ions with different m/z. To give a more detailed example, a false identification in a ^12^C^14^N (light) sample with a high light/heavy ratio, should not be reciprocally identified in the ^13^C^15^N (heavy) form, or if identified in the light form in the reciprocal experiment, it should not display an inverted low light/heavy ratio. If most of these cases reflect intrinsic experimental artefacts consistently present over biological replicates, independently of SILAC labeling swap, these false identifications should be prevalent in quadrants Q2 and Q4, because the peptide in question would be very unlikely misidentified in a reciprocal experiment due to its having a different m/z and/or display an inverted ratio. In such a context, consistency in quantitation over multiple biological replicates of label swapped experiments could be used as a parameter for efficiently excluding false identifications from final datasets, especially in the region of data points with high fold changes containing most of the key data that would be used for biological inference, such as for the identification of kinase substrates.

To test if performing a reciprocal labeling experiment indeed reduces false-positive identification and quantification, we estimated the error rate of phosphopeptide identifications by monitoring the distribution of reversed decoy hits from the list of phosphopeptide identifications that passed our basal quality criteria (PeptideProphet >0.9 and <20ppm precursor ion error). As shown in Figures 4A-B, decoy hits display a clear distribution bias towards quadrants Q2 and Q4, congruent with our rationale that false identifications are mostly unidirectional in quantitation and likely reflect artefacts that are extremely unlikely to occur in two reciprocal experiments, independently. Of all decoy hits in the unfiltered dataset, more than half (81 out of 130) were found to display ratios outside the 2-fold change range (Figs. 4A-B and 4E). Notably, we were able to remove all decoy hits from Q1 and Q3 (regions expected to contain key data for biological inference of true changes in phosphorylation events) using the Bow-tie filtering strategy in combination with a threshold of at least 2 PSMs per experiment (Figs 4C-E). Even when phosphopeptides reflected by 1 PSM per experiment were allowed in the dataset, the number of decoy hits in Q1 or Q3 remained low (2 hits) (Fig. 4D). These findings highlight the usefulness of our approach, which hinges on conducting a reciprocal SILAC experiment to improve confidence in both identification and quantitation in phosphoproteomic studies. Importantly, the described approach results in minor loss of valuable data content from low abundance phosphopeptides represented by only one PSM in each of the two reciprocal experiments.

**Figure 4.**
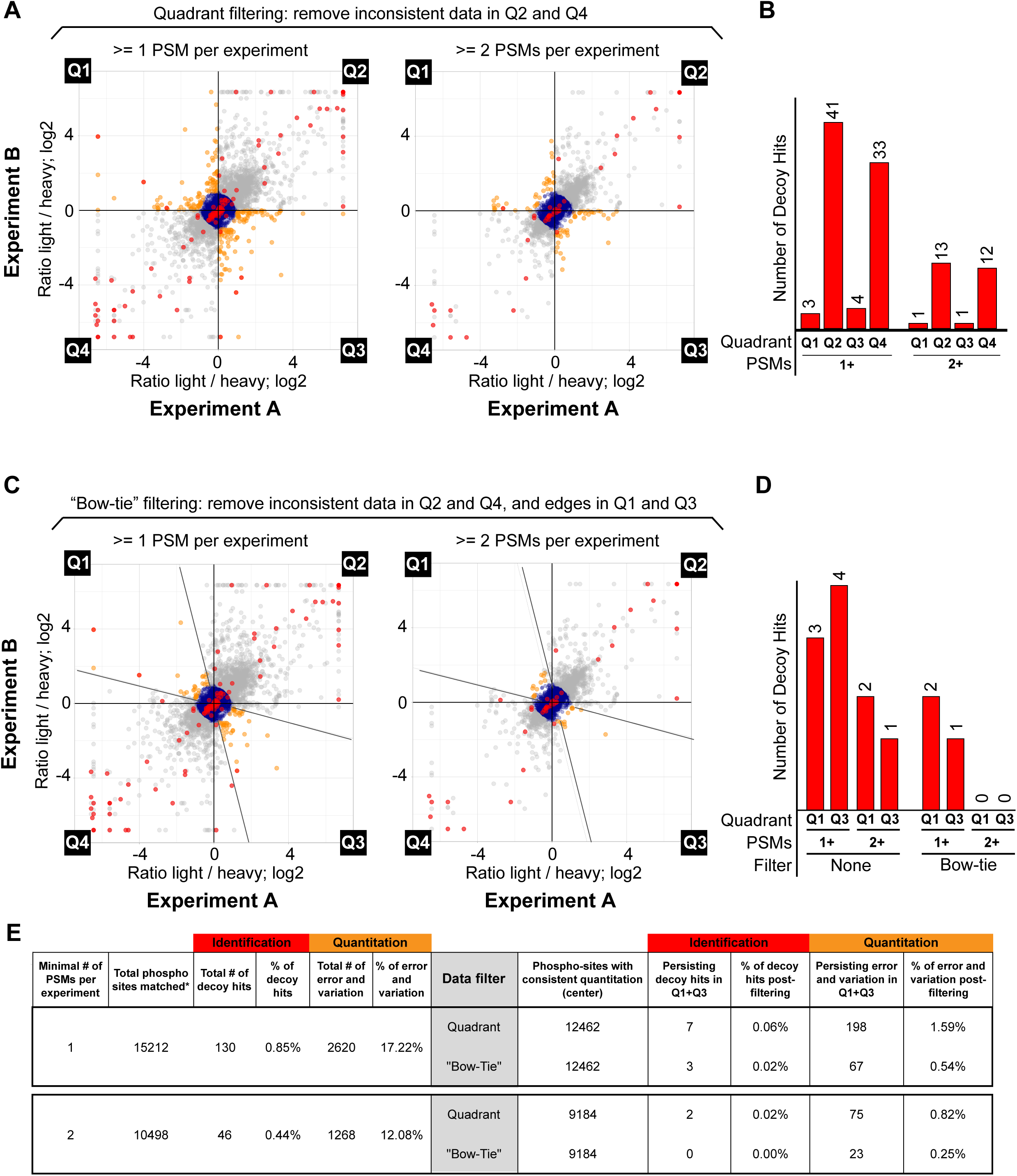
“Bow-Tie” Approach Efficiently Reduces Decoy Peptide Identifications. A. Scatterplot from Figure 2B. indicating hits from a decoy database. Decoy peptides are displayed in red. As in Figure 3, unfiltered EVs in Q1 and Q3 are displayed in orange. B. Histogram displaying number of decoy peptide hits in Q1 and Q3 with quadrant filtering applied. C. Scatterplot from Figure 2B. indicating hits from a decoy database and employment of Bow-tie filtering. Decoy peptides are displayed in red. As in Figure 3, unfiltered EVs in Q1 and Q3 after Bow-tie filtering are displayed in orange. D. Histogram displaying number of decoy peptide hits in Q1 and Q3 with Bow-tie filtering applied. E. Summary of phosphoproteomic data from Figure 2B comparing Quadrant to Bow-tie filtering, and 1 PSM cutoff to 2 PSM cutoff.

### Expanding the Mec1-dependent Signaling Network

We applied our optimized quantitative phosphoproteomic approach to the study of the DNA damage signaling kinase Mec1, the *Saccharomyces cerevisiae* ortholog of mammalian ATR. Mec1 is a phosphoinositide 3-kinase-related kinase (PIKK) kinase that controls of a range of nuclear processes [30]–[32]. We have previously used quantitative phosphoproteomics comparing WT and *mec1Δ* cells to uncover phosphorylation events dependent on Mec1 [13]. The results revealed an extensive network of Mec1 substrates, most of them phosphorylated at the preferential S/T-Q motif. Strikingly, application of our phosphoproteomic approach based on reciprocal SILAC swap and Bow-tie filtering dramatically expanded the network of Mec1-dependent phosphorylation events at the S/T-Q motif (Figs. 5A-C). We carried out the experiments in cells lacking the checkpoint adaptor Rad9 to minimize indirect downstream phosphorylation and preferentially reveal direct Mec1 substrates [33], [34]. Overall phosphoproteome coverage was similar to the control experiment, with approximately 20,000 phosphopeptide identifications for each SILAC reciprocal experiment. Application of our most relaxed filtering scheme, which considers phosphopeptides with 1 or more PSM in each experiment and a PeptideProphet score of 0.9 or greater, a total of 13,340 unique phosphopeptides from 2,774 different proteins were identified. The list of all phosphosites identified and regulated in our experiment is presented in Supplementary Table S2. As shown in Figure 5A, Q1 after Bow-tie filtering contained a large number of phosphosites consistently downregulated in cells lacking Mec1 in both reciprocal SILAC experiments. The number of phosphosites in Q1 was approximately equal to the number of EVs in Q2 and Q4 (Table S2), indicating that if experiments we performed using only one labeling scheme, many of these EVs excluded in our Bow-tie approach would have been erroneously called Mec1-dependent sites. Our filtering strategy allows minimal loss of data while increasing stringency for identification and exclusion of false positives through SILAC label swapping.

**Figure 5.**
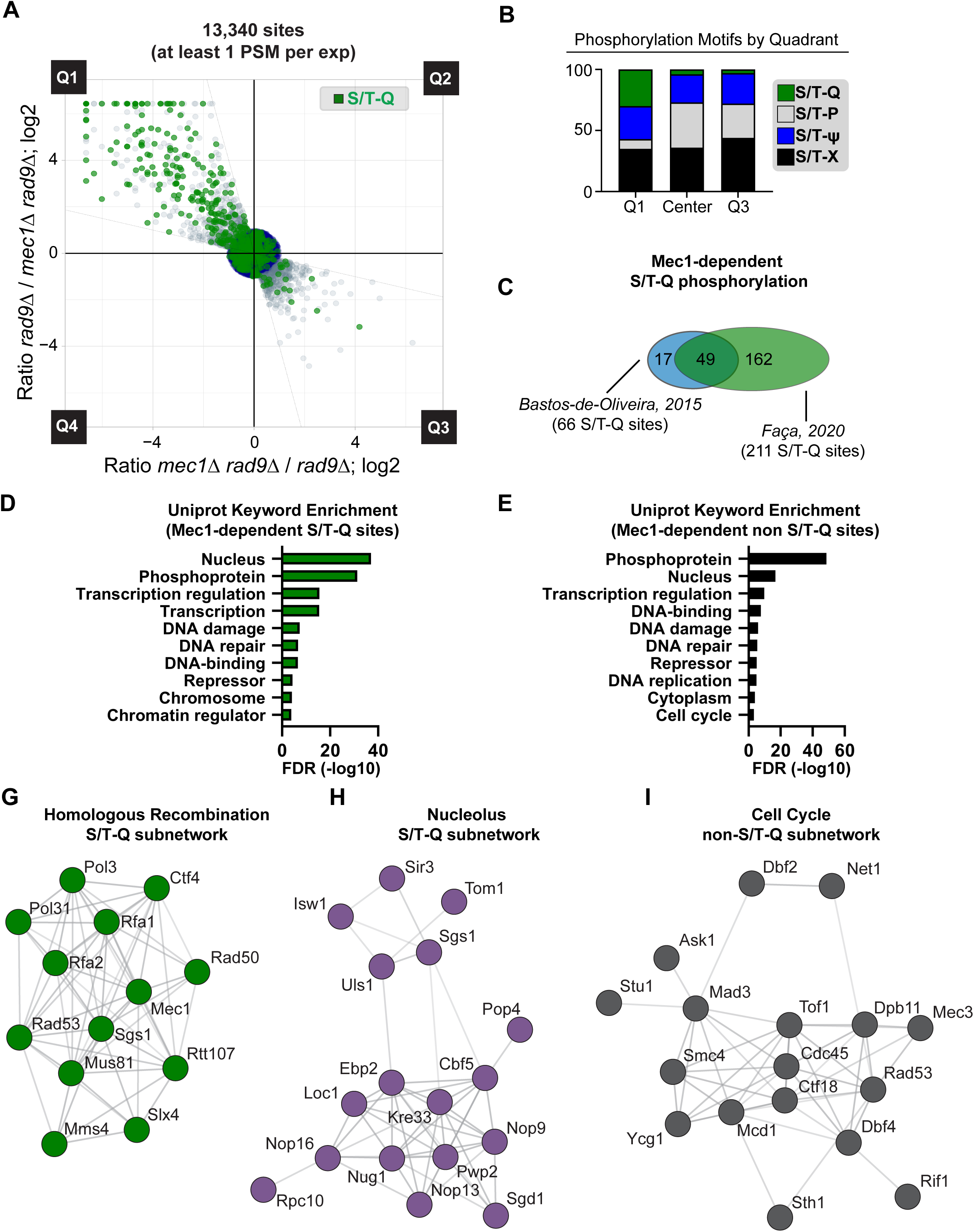
Expanding the Mec1 Signaling Network Using Low-PSM Threshold Coupled to “Bow-Tie” Filtering. A. Scatterplot (with Bow-tie filter applied; only data points within Bow-tie filter displayed) of forward and reciprocal SILAC experiment comparing phosphoproteome of *rad9*Δ cells to phosphoproteome of *rad9*Δ *mec1*Δ cells. Cells were treated with 0.02% MMS for 2hrs. B. Histogram depicting distribution of indicated phospho-motifs from data in Figure 5A. S/T-Q is the preferential motif for Mec1 phosphorylation. C. Venn-diagram of Mec1-dependent S/T-Q sites identified in this study compared to sites identified in our previous work [13]. D. Uniprot keyword enrichment analysis performed on all Mec1-dependent S/T-Q consensus phosphorylation sites from Figure 5A. E. Uniprot keyword enrichment analysis performed on all Mec1-dependent non-S/T-Q consensus phosphorylation sites from Figure 5A. F. String analysis of proteins with Mec1-dependent phosphorylation in the S/T-Q consensus revealed a sub-network of proteins involved in DNA repair via homologous recombination (HR). G. String analysis of proteins with Mec1-dependent phosphorylation in the S/T-Q consensus revealed a sub-network of proteins related to the nucleolus. H. String analysis of proteins with Mec1-dependent phosphorylation in non-S/T-Q consensus, revealed a sub-network of proteins related to cell cycle control.

To assess quality of the data generated using our SILAC-based filtering approach, and to retrieve meaningful biological insights about kinase action, we classified phosphosites according to their consensus motifs. Since Mec1 is known to preferentially phosphorylate the S/T-Q motif [13], [35], enrichment of this phospho-motif in Q1 would serve as an internal validation of the data. As shown in Figures 5A-B, Q1 is enriched with the preferential S/T-Q motif for Mec1 phosphorylation. Whereas S/T-Q motif represents only about 3% of the phospho-sites in the entire dataset, it represents 22% of the Mec1-dependent sites. Besides S/T-Q sites, Q1 also contained a number of sites with the S/T-Ψ (where Ψ denotes the bulky hydrophobic residues F, I, L and V) phospho motif (Fig. 5B), which is associated with the downstream checkpoint kinase Rad53 that is activated by Mec1 [36]. The occurrence of S/T-Ψ phosphorylation in the absence of the major *RAD9*-dependent pathway of Rad53 activation likely reflects Rad53 activation via the Mrc1 adaptor [37]. Indeed, Mec1-dependent phosphorylation sites were detected in Rad53, several of which are known Rad53 autophosphorylation sites [38] and indicate that this kinase is activated in *rad9Δ* cells expressing Mec1. We also identified Mec1-dependent phosphorylation sites in the Dun1 kinase, which is known to function downstream of Mec1 and Rad53 in the canonical DNA damage checkpoint signaling pathway [39]–[41]. Interestingly, our SILAC-based filtering approach revealed a number of Mec1-dependent sites that did not contain the S/T-Q or S/T-Ψ consensus, raising the possibility that Mec1 regulates the action of other kinases in addition to Rad53 and Dun1 in response to DNA damage.

In total, our quantitative phosphoproteomic approach using Bow-tie filtering of inconsistent ratios resulted in the identification of 211 S/T-Q Mec1-dependent phosphosites, which at least triples the number of Mec1 targets identified compared to our previous screen (Fig. 5C). Gene enrichment analysis of all Mec1-regulated S/T-Q sites (those with a log2 ratio >1 in WT cells relative to *mec1*Δ) based on their Uniprot annotation was consistent with our previous study showing that the substrate repertoire of this kinase was enriched for nuclear proteins involved in DNA repair, chromatin modification, and transcription (Fig. 5D). Analysis of Mec1-regulated sites containing a consensus motif that was not S/T-Q revealed that the scope of Mec1’s downstream signaling also largely encompassed proteins related to DNA damage, repair, and transcription. Interestingly, the non-S/TQ gene enrichment analysis showed a prevalence of cell-cycle, DNA replication, and cytoplasmic proteins (Fig. 5E). String network analysis [42] of the network of S/T-Q sites revealed extensive Mec1-dependent phosphorylation of components of the homologous recombination machinery (Fig. 5F), including proteins such as Rad50 that act early in HR during the resection step [43], [44], as well as proteins that act later during HR to regulate the processing of joint molecules, such as Sgs1 and Mus81-Mms4 [45]–[47]. Additionally, we found extensive Mec1-dependent phosphorylation of nucleolar proteins with the S/T-Q consensus (Fig. 5G). String analysis of non-S/T-Q substrates in the “cell cycle” node revealed non-canonical Mec1-dependent phosphorylation of the spindle assembly protein Mad3 and the condensing subunit Smc4 (Fig. 5H). Overall, these findings reveal the efficacy of our Bow-tie filtering method in identifying novel kinase substrates at high depth and specificity while minimizing false positives.

### In-Depth Phosphoproteomics Coupled to Bow-tie Filtering Uncovers the Tel1 Signaling Network

Tel1 is a DNA damage signaling protein kinase in the PIKK family closely related to Mec1 and orthologous to mammalian ataxia telangiectasia mutated (ATM) kinase [48], [49]. Like Mec1, Tel1 has been shown to phosphorylate proteins at the preferential S/T-Q motif [36], [50]. In yeast, a very limited number of Tel1 substrates have been reported, suggesting that the relative contribution of Tel1 to the overall DNA damage signaling response is smaller compared to Mec1. Indeed, in our previous screen for Tel1 targets we were only able to identify 4 phosphosites that are highly dependent on Tel1 [13]. We reasoned that the difficulties in delineating targets of Tel1 signaling in previous attempts were due to limitations in the coverage of phosphoproteomic analysis and issues in retrieving high confidence quantitative data for low abundant, underrepresented, phosphopeptides. To expand our knowledge of the Tel1 signaling network we used in depth phosphoproteomics coupled to Bow-tie filtering of EVs, comparing the phosphoproteome of wild-type and *tel1Δ* cells treated with the DNA alkylating drug MMS. In order to maximize coverage of the phosphoproteome, we performed a more extensive pre-fractionation of phosphopeptides using hydrophilic interaction liquid chromatography (HILIC). We analyzed a total of 32 fractions for each individual Tel1-dependency reciprocal experiment. In each individual experiment we detected around 30,000 phosphorylation sites. A total of 22,947 phosphorylation sites contained at least 1 PSM in each of the reciprocal experiments (Fig. 6A), including 10,883 phosphorylation sites not yet reported in the publicly available YeastMine repository that compiles most of the phosphorylation sites reported for budding yeast (21,335 sites as in November of 2019) (Balakrishnan et al., 2012). Upon application of Bow-tie filtering, a total of 17,362 sites remained within the constraints of our filter. Strikingly, 416 regulated phosphorylation sites were present in quadrant Q1, indicating a large number of sites dependent on the Tel1 kinase (Fig. 6A). The results reveal an enrichment for the S/T-Q motif in Q1, defining a total of 68 Tel1-dependent S/T-Q sites (Fig. 6B). This novel set of Tel1-dependent S/T-Q sites represents a drastic increase compared to the 4 Tel1-dependent S/T-Q sites detected in our previously published dataset (Fig. 6C). Unexpectedly, we also observed strong enrichment of phospho-motifs with an acidic residue (D or E) at the -1 position of Tel1-dependent phosphorylation sites (Figs. 6A and 6B), suggesting a novel preferential phospho-motif for Tel1 phosphorylation or that Tel1 may activate a downstream kinase that preferentially targets D/E-S/T motif. We favor the former possibility, since 42 of the 68 Tel1-dependent S/T-Q sites also contain a D or E at the -1 position. Since many of the Tel1-dependent D/E-S/T sites do not contain a Q at the +1 position, Tel1 may also directly phosphorylate non-S/T-Q motifs, as long as there is a D or E at the -1 position. However, this model does not exclude the possibility of an alternative downstream kinase also targeting some, or most, of the D/E-S/T sites. Indeed, we detected a Tel1-dependent glutamine-directed phosphorylation in the Hrr25 kinase (serine 438), which is a CK1-like kinase known to prefer acidophilic motifs [51]. Analysis of Uniprot annotations enrichment of Tel1-dependent S/T-Q sites showed that, as expected, most Tel1 phosphosubstrates are related to nuclear processes, DNA damage, and DNA repair (Fig. 6D). String network analysis of the DNA damage/repair subnetwork of Tel1 phosphosubstrates showed that many of the proteins in this subcategory share genetic or physical interactions (Fig. 6E), although Hrr25 does not interact with any of the other repair proteins we identified as being Tel1-dependent. Analysis of D/E-S/T consensus Tel1-dependent phosphorylation revealed a similar preference of Tel1 for nuclear substrates involved in DNA damage and repair (Fig. 6F). Unexpectedly, Tel1-dependent phosphorylation of a number of RNA Polymerase II holoenzyme subunits bearing the D/E-S/T consensus was observed, raising the possibility that Tel1 regulates RNAPII function (Fig. 6G).

**Figure 6.**
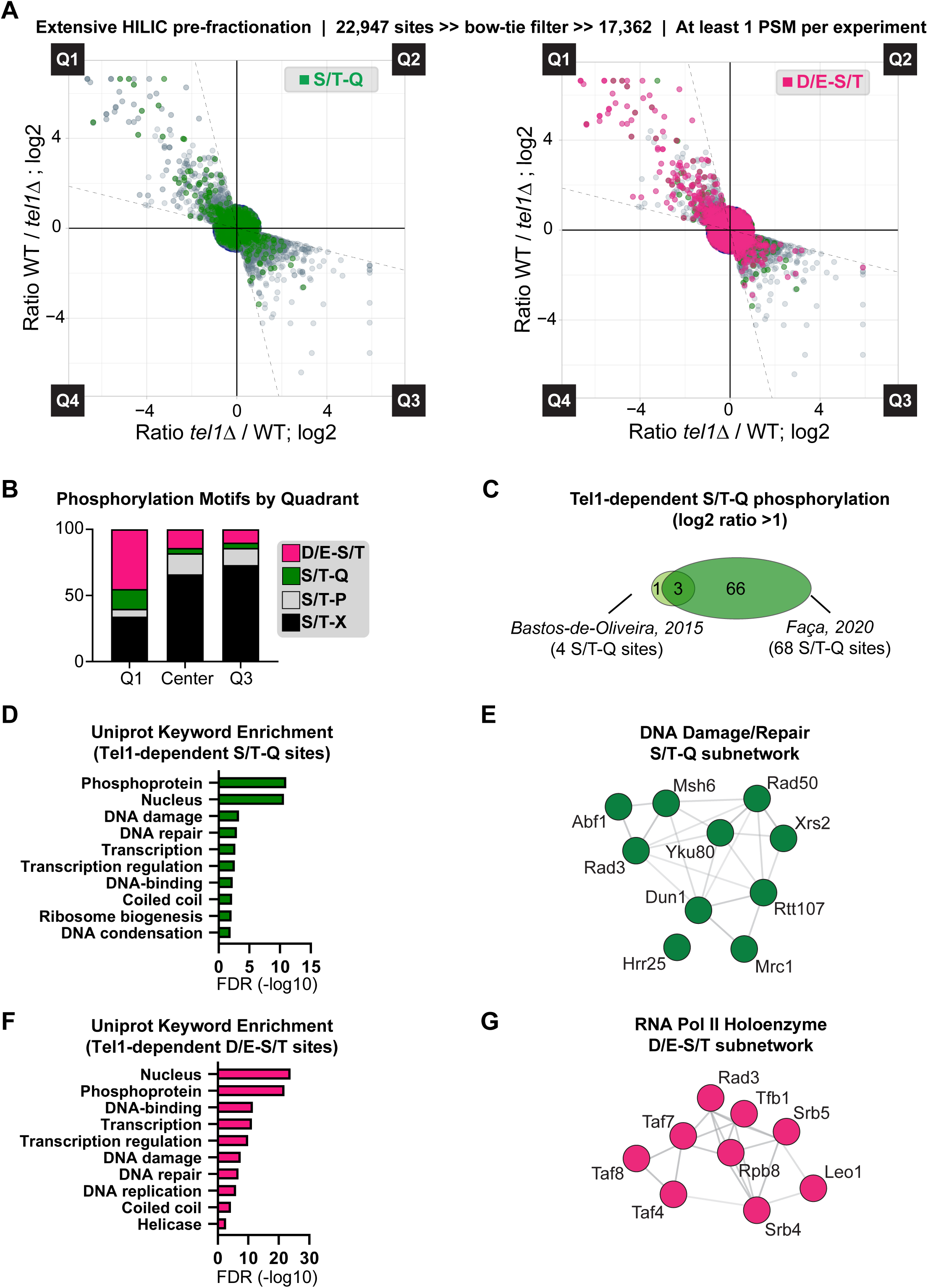
Expanding the Tel1 Signaling Network Using Extensive HILIC Pre-fractionation and Low-PSM Threshold Coupled to “Bow-Tie” Filtering. A. Scatterplots (with Bow-tie filter applied; only points within Bow-tie filter displayed) of forward and reciprocal SILAC experiment comparing the phosphoproteomes of wild-type and *tel1*Δ cells. S/T-Q motifs highlighted in green (left). D/E-S/T motifs highlighted in pink (right). Cells treated with 0.1% MMS for 2hrs. B. Histogram depicting distribution of indicated phospho-motifs from data in Figure 6A. C. Venn-diagram of Tel1-dependent S/T-Q sites identified in this study compared to sites identified in our previous work [13]. D. Uniprot keyword enrichment analysis performed on all Tel1-dependent S/T-Q consensus phosphorylation sites from Figure 6A. E. String analysis of proteins with Tel1-dependent phosphorylation in the S/T-Q consensus revealed a sub-network of proteins related to DNA damage and repair. F. Uniprot keyword enrichment analysis performed on all Tel1-dependent D/E-S/T consensus phosphorylation sites from Figure 6A. G. String analysis of proteins with Tel1-dependent phosphorylation in the D/E-S/T consensus revealed a sub-network of proteins of the RNA Polymerase II holoenzyme.

These results considerably expand our knowledge of the Tel1 signaling network. The use of Bow-tie filtering was crucial for deriving high quality quantitative data from phosphopeptides represented by very few PSMs, which allowed us to dig deeper into the dataset. For example, several phosphopeptides from proteins involved in the Tel1-regulated pathways, such as Rif1 [52], [53] and Rad50 [54], [55], were only detected once in each of the reciprocal experiments. Overall, the findings establish that quantitative analysis of tens of thousands of phosphosites can be achieved with robustness and high confidence.

## DISCUSSION

The field of phosphoproteomics has made significant strides toward improved phosphopeptide detection and quantitation since the seminal paper by Fenselau et al, which described the first application of FAB mass spectrometry for phosphopeptide characterization [56]. Throughput as well as robustness has increased, and modern instruments and workflows can routinely detect and quantitate thousands of phosphopeptides in a single run. Still, the intrinsic issue of lack of redundancy in data representation for each phosphopeptide remains, leading to lack of statistical power for generating high confidence quantitation and identification for large portions of the dataset, especially for low abundance phosphopeptides that often rely on single PSMs with noisy signals. This issue has been tackled predominantly by requiring higher numbers of PSMs per phosphopeptide, with the consequent trade-off of eliminating a substantial fraction of the dataset that may contain most of the biologically meaningful regulatory, and low abundant, events. In this work we present a workaround that allows for the efficient exclusion of technical noise and variation through the use of a reciprocal SILAC experiment, while allowing for the identification and quantitation of low abundance phosphopeptides. We leveraged the sensitivity of this new pipeline by combining the proposed Bow-tie analysis with samples that had been extensively prefractionated using HILIC chromatography. The result is a drastic expansion in coverage with concomitant reduction in error and technical variation in the overall quantitative data. This combination of high specificity, low-PSM phosphoproteomics with HILIC, which is particularly suited to phosphopeptide fractionation due to the hydrophilicity of the phosphate group [57] revealed ∼15,000 unique phosphopeptides in a short fractionation schema (15 fractions) and ∼20,000 in longer gradients (32 fractions), as demonstrated for our identification of Tel1 targets.

Central to the Bow-tie filtering strategy presented in this study is the use of metabolic labelling with stable isotopes (SILAC) and the consequent shift in mass of parent and fragment ions of phosphopeptides. Such a large delta mass between phosphopeptides in reciprocal experiments forces a stringent requirement in which phosphopeptide identification with inverted fold change in each experiments should also exhibit proper delta mass shift. In addition to allowing efficient detection of EVs, this approach also led to a dramatic reduction in the number of decoy database peptide identifications in quadrants 1 and 3. This serves as direct proof that reciprocal labeling reduces false-positive identification and associated quantitation. False-positives identifications as proposed to represent either artefacts, exogenous sample contaminants not represented in the searched database, or contain other types of modifications not considered in our search as variable modifications [58]. The Bow-tie approach applied to the mock dataset reduced false-positive hits to virtually zero, when Q1 and Q3 were considered and at a PeptideProphet score minimum score of 0.9, satisfying our needs for highly sensitive and comprehensive strategy to uncover phosphopeptides of low abundance and low PSM counts. A near-zero frequency of false-positive identifications appearing in Q1 and Q3 is essential to our SILAC-based approach, because peptide identification essentially serves as the most important “gatekeeper” of filtering meaningful biological data from technical noise.

The mass spectrometric data processing pipeline employed in this study relied on the Trans-Proteomic Pipeline (TPP) suit of proteomic tools, including updated tools for peptide identification with COMET [27] and scoring with PeptideProphet [28]. For SILAC quantitation, we used the Xpress precursor ion intensity quantitation tool [59], and for phosphosite localization and scoring we used the newly described PTMProphet tool [29]. PTMProphet models the potential sites of phosphorylation independently of the spectrum identification provided by the search engine and calculates probabilities for each potential modification site. This feature allowed us to design additional steps in our R-based scripts for handling clustered S/T/Y residues, which is a common occurrence in phosphopeptides. Unambiguous, high-confidence phosphorylated S/T/Y residues with neighboring S/T/Y residues were kept separate; medium or low-confidence phosphorylated S/T/Y residues with adjacent S/T/Y residues were combined with their neighbors and considered in our subsequent analyses as a “cluster.” In our filtered data for Tel1-dependent screen, 2072 (approximately 12%) of these phosphopeptides were observed. A non-systematic comparison of other combinations of data processing and site localization tools suggested that the chosen TPP-based set of tools was the most sensitive and efficient for our applications.

The ability of our Bow-tie approach to separate biologically meaningful phosphorylation from technical noise is exemplified by the observed regulation in our Mec1 phospho-mapping dataset. In contrast to the control dataset, in which there were a small number of points in quadrants 1 and 3, there were a number of regulated sites in Q1 in our Mec1 dataset (and much less in Q3). These regulated points in Q1, expectedly, were largely phosphosites with the S/T-Q consensus, indicating primary Mec1-dependent phosphorylation in response to DNA damage that was ablated in the absence of the *MEC1* gene. In addition to revealing many known Mec1 targets identified in other studies, which was our intention as a validation of our method, we revealed a number of previously unreported substrates of Mec1, including a number of proteins associated with the nucleolus. For example, we identified phosphorylation on serine 1007 (an S/T-Q site) of Kre33, a relatively understudied protein that promotes maturation of 18S rRNA [60], [61]. Future work should be targeted toward understanding how Mec1 signaling contributes to nuclear homeostasis independently of its established roles in activation of the DNA damage checkpoint.

As a final test of our Bow-tie pipeline, we defined a comprehensive list of Tel1 phosphosubstrates upon DNA damage which contained more than 15x the number of Tel1-dependent S/T-Q sites from our previous study [13]. Importantly, this experiment highlighted the increased depth and coverage achievable by our pipeline. This feature allowed us to not only observe many new Tel1-dependent S/T-Q sites, but also revealed an abundance of Tel1-dependent sites that contained the amino acids D or E at the -1 position. There are two possible explanation for this observed effect. One explanation is that Tel1 itself is acidophilic, and the presence of D/E amino acids at the -1 position directs phosphorylation of S/T-Q sites by Tel1. Another possible explanation is that Tel1 activates an additional kinase which contributes to the phosphorylation of the large number of non-S/T-Q sites we observe to be Tel1 dependent. A possibility raised by our data is that Tel1 may regulate the casein kinase homolog Hrr25, as we did observe a Tel1-dependent S438Q site in this protein. While most kinases are basophilic, the casein kinases are famously acidophilic in nature [51]. Tel1 activation of Hrr25 by an as-yet unknown mechanism could explain why we observe a large number of Tel1-dependent D/E-S/T phosphorylation sites.

Tel1 is crucial for telomere control maintenance [48], [49], [62], but the mechanism(s) by which Tel1 controls telomeres remains unknown. Our datasets contain a range of substrates that may represent the mechanistic link between Tel1 and telomere regulation. For example, we identified phosphorylation on S256 (consensus D/E-S/T-Q) of the ARS-associating factor Abf1 [63], a site not yet reported to be targeted by Tel1. Abf1 is both a transcriptional activator and a repressor, and it is functionally similar to the known telomere maintenance protein Rap1 [64]–[67]. Future yeast genetic work should be targeted at elucidating the exact mechanisms of Tel1 regulation of its many substrates and defining, mechanistically, how Tel1 controls telomere length regulation.

In summary, here we report a simple and robust SILAC-based phosphoproteomic data analysis pipeline that allows for identification and quantitation of phosphopeptides with high confidence and coverage. The depth of the analyses allowed identification of a range of novel substrates of both the kinases Mec1 and Tel1, including the Hrr25 kinase as a potential downstream kinase regulated by Tel1. While this work highlights the utility of SILAC for high confidence and in depth quantitative phosphoproteomics, the same rationale could be applied to improve the quantitative analysis of other low-abundance post-translational modifications such as sumoylation, ubiquitylation, and acetylation.

## ACKNOWLEDGEMENTS

We thank Beatriz S. Almeida for technical support; we thank all members of the Smolka Lab for valuable discussions related to this work. This work is supported by grants from the National Institute of Health (R01-GM097272, R01-HD095296 and R01-GM123018) to M.B.S.

## PROTEOMICS DATA AVAILABILITY

Mass spectrometry data generated from this study has been deposited to the Massive database (http://massive.ucsd.edu). The control mock experiment data received the ID: MSV000084852, doi:10.25345/C58M3B, and ProteomeExchange ID: PXD017322. The Mec1 targets experiment data received the ID: MSV000084875, doi:10.25345/C56Q44, and ProteomeExchange ID: PXD017339. The Tel1 targets experiment data received the ID: MSV000084871; doi: 10.25345/C5KX2V and ProteomeExchange ID: PXD017338.

## AUTHOR CONTRIBUTIONS

VMF and MBS conceptualized the project and the pipeline. EJS and WJC performed experiments. All authors contributed to data analysis. JT and VMF wrote the data analysis script. EJS, VMF, and MBS wrote the paper.

